# A *Plasmodium falciparum* genetic cross reveals the contributions of *pfcrt* and *plasmepsin II/III* to piperaquine drug resistance

**DOI:** 10.1101/2023.06.06.543862

**Authors:** John Kane, Xue Li, Sudhir Kumar, Katrina A. Button-Simons, Katelyn M. Vendrely Brenneman, Haley Dahlhoff, Mackenzie A.C. Sievert, Lisa A. Checkley, Douglas A. Shoue, Puspendra P. Singh, Biley A. Abatiyow, Meseret T. Haile, Shalini Nair, Ann Reyes, Rupam Tripura, Tom Peto, Dysoley Lek, Stefan H.I. Kappe, Mehul Dhorda, Standwell C Nkhoma, Ian H. Cheeseman, Ashley M. Vaughan, Timothy J. C. Anderson, Michael T. Ferdig

**Author notes:** Address correspondence to Michael T. Ferdig. Present addresses: Katelyn Vendrely Brenneman, Harvard T.H. Chan School of Public Health, Boston, MA, USA; Puspendra P. Singh, ECD Division, Indian Council of Medical Research, New Delhi, India.

## Abstract

Piperaquine (PPQ) is widely used in combination with dihydroartemisinin (DHA) as a first-line treatment against malaria parasites. Multiple genetic drivers of PPQ resistance have been reported, including mutations in the *Plasmodium falciparum chloroquine resistance transporter* (*pfcrt*) and increased copies of *plasmepsin II/III* (*pm2/3*). We generated a cross between a Cambodia-derived multi-drug resistant KEL1/PLA1 lineage isolate (KH004) and a drug susceptible parasite isolated in Malawi (Mal31). Mal31 harbors a wild-type (3D7-like*) pfcrt* allele and a single copy of *pm2/3,* while KH004 has a chloroquine-resistant (Dd2-like*) pfcrt* allele with an additional G367C substitution and four copies of *pm2/3*. We recovered 104 unique recombinant progeny and examined a targeted set of progeny representing all possible combinations of variants at *pfcrt and pm2/3* for detailed analysis of competitive fitness and a range of PPQ susceptibility phenotypes, including PPQ survival assay (PSA), area under the dose-response curve (AUC), and a limited point IC_50_ (LP-IC_50_). We find that inheritance of the KH004 *pfcrt* allele is required for PPQ resistance, whereas copy number variation in *pm2/3* further enhances resistance but does not confer resistance in the absence of PPQ-R-associated mutations in *pfcrt*. Deeper investigation of genotype-phenotype relationships demonstrates that progeny clones from experimental crosses can be used to understand the relative contributions *of pfcrt, pm2/3,* and parasite genetic background, to a range of PPQ-related traits and confirm the critical role of the PfCRT G367C substitution in PPQ resistance.

**Importance:** Resistance to PPQ used in combination with DHA has emerged in Cambodia and threatens to spread to other malaria-endemic regions. Understanding the causal mutations of drug resistance and their impact on parasite fitness is critical for surveillance and intervention, and can also reveal new avenues to limiting the evolution and spread of drug resistance. An experimental genetic cross is a powerful tool for pinpointing the genetic determinants of key drug resistance and fitness phenotypes and have the distinct advantage of assaying the effects of naturally evolved genetic variation. Our study was significantly strengthened because the full a range of copies of KH004 *pm2/3* was inherited among the progeny clones, allowing us to directly test the role *of pm2/3* copy number on resistance-related phenotypes in the context of a unique *pfcrt* allele. Our multi-gene model suggests an important role for both loci in the evolution of this ACT resistant parasite lineage.

## Introduction

Malaria is a life-threatening parasitic disease that puts nearly half of the world’s population at risk (1). The World Health Organization estimates that in 2020 there were 247 million malaria cases and 619,000 deaths reported across 84 countries, with 76% of those deaths occurring in children under the age of 5 (1). The continued evolution of drug-resistant *Plasmodium falciparum* asexual blood-stage parasites has been one of the major obstacles to global malaria elimination and eradication efforts (1). Current frontline treatments for uncomplicated *P. falciparum* infection are artemisinin-based combination therapies (ACT), which are composed of a fast-acting artemisinin (ART) derivative paired with one or more longer-lasting partner drugs (2). ACTs began to replace historically efficient antimalarials, such as chloroquine (CQ) and sulfadoxine-pyrimethamine (SP), as first-line therapies in the early 2000s after resistance to these previous therapies swept throughout Africa (3–5). The WHO has recommended ACTs for uncomplicated malaria since 2006 due to their greater efficacy and decreased resistance compared to other single-drug therapeutics, with the most common ACTs used in Southeast Asia (SEA) being dihydroartemisinin/piperaquine (DHA+PPQ), artemether/lumefantrine (AL), and artesunate/mefloquine (AS+MQ). The ACT combination DHA-PPQ has been widely used throughout the Greater Mekong Subregion (GMS) of SEA, which has historically been a hotbed for the evolution of antimalarial drug resistance (6–8).

The spread of resistance to DHA+PPQ has been attributed to the rapidly expanding, multidrug-resistant KEL1/PLA1 lineage of parasites (9–11). These parasites originally emerged out of western Cambodia in 2008 and have since spread to Vietnam, Laos, and northeastern Thailand (10). KEL1/PLA1 parasites are characterized by ART-resistant mutations in *kelch13* (KEL1), specifically the C580Y substitution, as well as a copy number (CN) amplification of the two aspartic protease genes *plasmepsin II/III* (PLA1) associated with decreased susceptibility to PPQ (9, 10, 12, 13). Reduced susceptibility to ART component of and ACT results in a greater remaining parasite load that has to be cleared by the partner drug, increasing the selective pressure for ACT failure (9, 14–17).

After the original detection of PPQ resistance in Cambodia in 2015, independent genome-wide association studies identified an association between *plasmepsin II/III* (*pm2/3*) variants and PPQ-R (16, 18–20). The mechanism proposed to support this association is an amplification of *pm2/3* that reduces concentrations of reactive heme in the parasite digestive vacuole (DV) to counter the inhibitory action of PPQ (20). Increased parasite survival at higher PPQ concentrations amongst Cambodian field isolates was associated with the amplification of *pm2/3* (13). Further *in vitro* studies of *pm2/3* amplification found that in the PPQ-S 3D7 background, inactivation of *pm2/3* results in a mild increase in PPQ susceptibility, whereas overexpression of *pm2/3* alone did not impact the degree of parasite susceptibility to PPQ, artesunate (AS), or CQ (21, 22). Nevertheless, while *pm2/3* was initially heralded as a defining feature of PPQ-R, variations of *in vitro* and isolate-based genotype-phenotype associations suggest that additional loci are needed to fully account for resistance (13, 23).

Beyond the role of *pm2/3*, decreased copies of *pfmdr1* and single nucleotide polymorphisms (SNP) in several other genes, including *exonuclease* (PF3D7_1362500) and the *P. falciparum chloroquine resistance transporter* (*pfcrt*) have been associated with PPQ-R parasites (16, 18, 24–26). Novel haplotypes containing amino acid substitutions in PfCRT were initially identified due to their increasing prevalence in areas utilizing DHA+PPQ, and *in vitro* studies showed that isogenic parasites edited with these substitutions alone can impact parasite uptake of PPQ and drug susceptibility independent of *pm2/3* copy number (25, 27–30). Field studies in the GMS report a continued expansion of some of these *pfcrt* haplotypes in the last five years that often co-occur in parasites carrying multiple copies of *pm2/3*, suggesting there is no fitness cost associated with this allelic combination (31, 32). Well studied PfCRT substitutions linked to PPQ-R, including H97Y, F145I, M343L, G350R, and G353V, arose in either a Dd2- or 7G8-like CQ-resistant (CQ-R) genetic background (33, 34).

Compared to other antimalarial drugs, assessing *in vitro* PPQ-R has been challenging due to the unusual dose-response effect in PPQ-R parasites. These resistant parasites show incomplete killing at the higher concentrations, resulting in an unusual bimodal dose-response curve (13). To account for this non-traditional drug concentration-biological effect relationships, various modifications to standard IC_50_ values have been used, including total area under the curve (AUC) (13). Alternatively, the piperaquine survival assay (PSA) was developed to measure the parasite survival rate after exposure to a single pharmacologically relevant (200 nM) dose of PPQ (18). While these different readouts will correlate in their characterization of resistance, they also likely capture distinct aspects of the drug-parasite interaction. We predict that genetic loci will contribute differentially to these different response readouts, perhaps pointing to distinct underlying biological in progeny inheriting unique allele combinations. .

The genetic background of *P. falciparum* has been shown to play a central role in the emergence of drug resistance and its impact on parasite fitness (35–39). While several individual genes have been implicated in PPQ-R, the understanding how these genes interact is lacking any may provide insights about how parasites develop resistance and how some lineages have strong potential to expand. To further dissect the contributions of different loci to PPQ-R, here we examined progeny from a genetic cross between a multidrug-resistant Cambodian parasite of KEL1/PLA1 lineage (KH004) and a drug-sensitive Malawian isolate (Mal31). KH004 carries a C508Y Kelch13 substitution associated with ART-R, a Dd2-like PfCRT (74I, 75E, 76T, 220S, 271E, 326S, 356T, and 371I), associated with CQ-R, with an additional G367C substitution, and 4 copies of *pm2/3*. Mal31 has a wild-type genotype at *kelch13* and *pfcrt* and carries a single copy of *pm2/3* (Table S1). The G367C substitution in PfCRT carried by KH004 has previously been identified as a novel haplotype present in SEA. However, it’s association with *in* vitro PPQ resistance has not been determined (25, 33, 40).

Pairing a novel genetic cross with quantitative trait loci (QTL) mapping can be a powerful tool for quantifying the contributions of all loci to phenotypes of interest. We take a multi-phenotype linkage approach to validate and characterize known resistance determinants as well as identify novel secondary loci associated with each unique resistance-related phenotype, underscoring slight differences in biological features of these measured traits. We find that increasing levels of resistance to PPQ prominently impacted by *pfcrt,* specifically a novel G367C substitution, with additional contributions from *pm2/3* amplification and other novel loci.

## Results

### Experimental genetic crosses

We generated two biological replicate genetic crosses between Mal31 and KH004 using *Anopheles stephensi* mosquitoes and human liver-chimeric FRG huHep mice, as described in Vaughan *et al.* (41). There are 14,455 core-genome SNPs (defined in Miles et al. (42)) distinguishing these two parental parasites (43) and both parental parasites were cloned by limiting dilution. We generated the two recombinant pools for this cross using independent groups of 250 mosquitoes and intravenously injected salivary gland sporozoites from these two pools into separate FRG huHep mice. The oocyst prevalence from the two pools were 21% and 33% respectively and the average number of oocysts, 0.5 and 0.6. We were able to isolate approximately 225,000 salivary gland sporozoites from each pool, approximately 900 sporozoites per mosquito. Results from the parental feeds alone also showed a low oocyst prevalence (Mal31, 41% and KH004, 33%, with total sporozoite numbers per mosquito averaging 800 for Mal31 and 480 for KH004. The total number of potential recombinants from each pool (assuming 50% of oocysts are recombinant and 50% are parental), was 0.5 oocysts × 4 recombinants per oocyst x 250 mosquitoes x 0.5 = 250 for pool 1 and 0.6 oocysts x 4 recombinants × 250 mosquitoes x 0.5 = 300 for pool 2. The initial allele frequencies of Mal31 of the two recombinant pools directly from mice were 0.70 and 0.66, suggesting the existence of Mal31 selfed progeny.

We cloned and sequenced 567 progeny from the two recombinant pools (286 from pool 1 and 281 from pool 2) (Table S2). Initial analysis led to the removal of 153 non-clonal cultures (F_WS_ <0.9, or non-clonal based on allele frequency plot). Of the 414 clonal progeny cultures, 92 were selfed Mal31, 3 were selfed KH004, and 319 were recombinant. We identified a total of 104 unique recombinants (identical by descent [IBD] clusters), of which 65 were from pool 1 and 39 were from pool 2. There were 44 IBD clusters with more than 2 clones, and 60 were singletons. The results show a skewing towards selfing of Mal31, as predicted and overall, 319/414 = 77% of the cloned progeny were recombinant.

### Determination of plasmepsin II/III copy numbers in parental parasites and progeny

We measured *pm2/3* CN for parental parasites with both nanopore long reads and Illumina short reads (Figure 1A). Mal31 harbors one copy of *pm2/3*, while KH004 harbors multiple copies of *pm2/3.* We were able to extract four long reads from KH004 nanopore sequences that cover the *pm2/3* genes alongside >10kb flanking regions (both upstream and downstream) and two of the reads had 5 copies, one had 4, and one had 3 copies of *pm2/3*.

**Figure 1:**
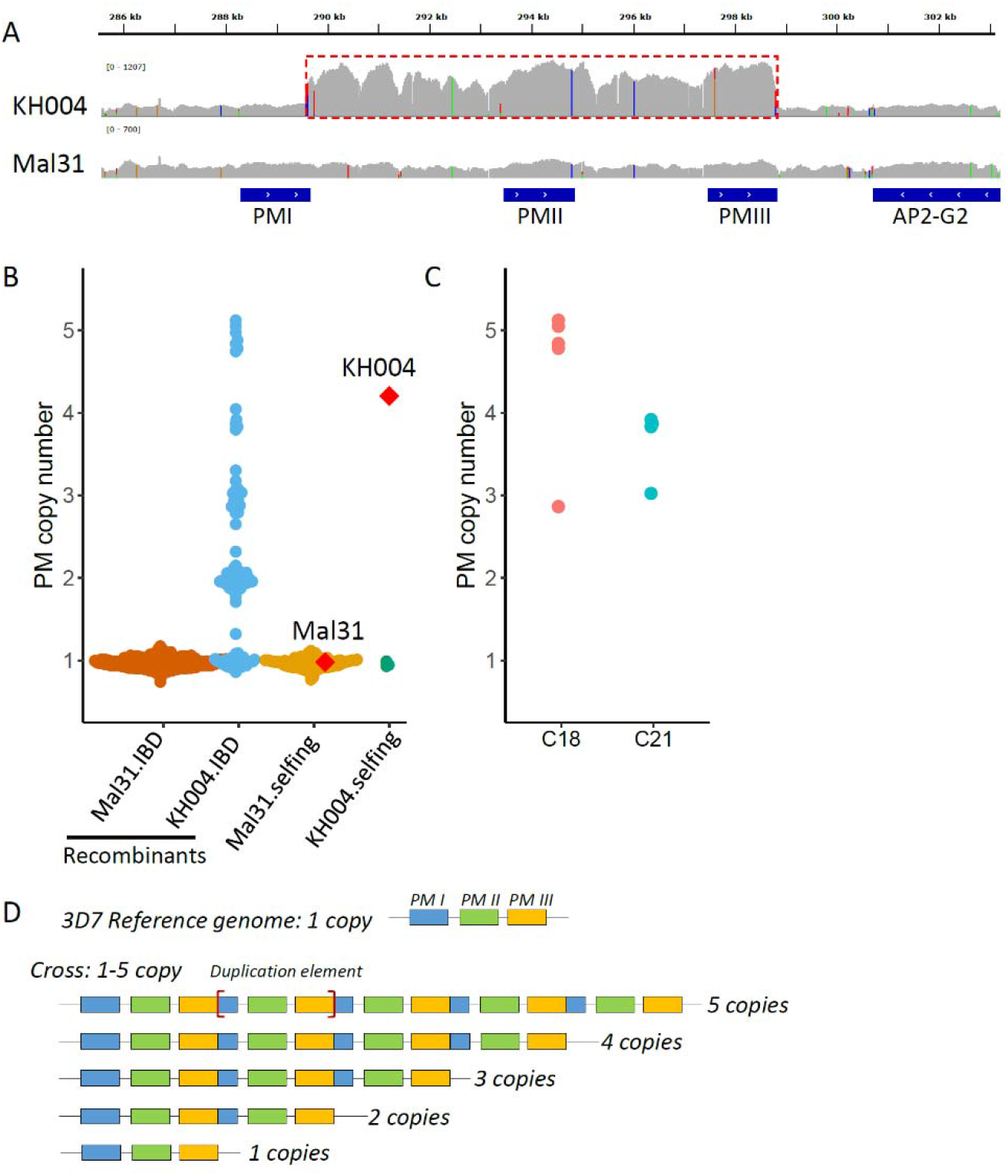
**Plasmepsin II/III (pm 2/3) copy number variations in parents and progeny**. A) IGV plot showing copy number variations in parental parasites at pm gene region with the pm repeat unit labeled with a red box. B) pm 2/3 copy number of parental parasites and recombinant and selfed progeny. Mal31.IBD indicates recombinant progeny with pm allele inherited from Mal31; KH004.IBD represents progeny with pm from KH004. Parental parasites Mal31 and KH004 are labeled with red diamonds. C) rapid evolution of pm 2/3 copy numbers in identical-by-descent (IBD) progeny. C18 and C21 are two IBD progeny groups. Of which, C18 contains 5 clones, with 4 of them containing 5 copies and 1 of them containing 3 copies of pm; while C21 contains 4 clones, with 3 of them contain 4 copies, and 1 contains 3 copies of pm. D) Nanopore long reads that cover the pm 2/3 CNV and flanking regions. The boundaries of the repeat unit are labeled with [red square brackets].

We further used Illumina short reads to analyze *pm2/3* CN in progeny (Figure 1B and C) which revealed that *pm2/3* CN ranges from 1 to 5 in progeny; all selfed KH004 contained only one copy of *pm2/3*, in contrast to the parental KH004 that had multiple copy of *pm2/3*; copy number variations (CNV) of *pm2/3* were detected inside IBD clusters, i.e., C18 and C21 (Figure 1C), suggesting rapid CNV evolution during *in vitro* blood culture and/or meiosis.

### Assessment of novel allelic combinations of pfcrt and pm2/3

From our set of 104 unique recombinant progeny, parasites were grouped into 10 unique clusters based on the inheritance of *pfcrt* and *pm2/3* alleles as well as *pm2/3* CN (Table S3). Progeny carrying the Mal31 *pfcrt* and KH004 *pm2/3* alleles inherited all potential copies of *pm2/3*, from 1 copy to 5 copies, whereas no progeny carry KH004 *pfcrt* with more than 3 copies of *pm2/3.* Regardless of *pfcrt* inheritance, only 28% (13/46) of parasites which inherited the KH004 *pm2/3* allele have >2 copies, indicating a potential fitness cost to carrying an excess number of copies.

Genome-wide, we observed two large skews towards Mal31 alleles, one on chromosome (chr) 7, and the second on chr 14 (Figure S1). The skewed region on chr 7 is centered with *pfcrt*, which is known to carry high fitness cost with CQ resistance alleles (8). The apicoplast ribosomal protein S10 (ARPS10, PF3D7_1460900) is located at the peak of chr 14 skewed region. Selection against the same *ARPS10* allele (Val127Met and Asp128His) has been previously detected in two other independent crosses between Asian and African parasites (38, 44), indicating a strong fitness cost carried by this allele from a Southeast Asia parent (45).

We detected no skewed inheritance at *pm2/3*. We also saw no evidence for pairwise linkage disequilibrium or co-inheritance between *pfcrt, kelch13*, or *pm2/3* alleles. However, we did observe a significant (p = 0.006) association between inheritance of the KH004 ART-R *kelch13* allele and multiple copies of *pm2/3* (Table S4).

### PfCRT mutations are the major contributor to PPQ resistance

A major challenge to quantifying response to PPQ is the biphasic dose-response curve which suggests incomplete parasite killing at higher drug concentrations as has been previously reported in PPQ-R parasites (13). To comprehensively assess the impact of novel combinations of *pfcrt* and *pm2/3* on parasite response to PPQ, we measured three unique phenotypes - PSA, AUC, and a modified IC_50_ based on a limited-point dose response curve (LP-IC_50_) in both parents from the KH004 × Mal31 cross as well as 48 select progeny chosen to encompass all available combinations of *pfcrt* and *pm2/3* CN.

IC_50_ is not traditionally used for phenotyping PPQ is due to the atypical biphasic drug response curve generated by resistant parasites (13).We find that progeny parasites that inherited the KH004 *pfcrt* allele and also multiple copies of *pm2/3* generate a biphasic dose response curve, whereas progeny with only one copy of *pm2/3* and carrying the Mal31 *pfcrt* allele produce a standard sigmoidal dose response (Figure S2A). This observed range of dose- response curve shapes within our progeny set affirms our use of multiple trait measurements, including a modification of standard IC_50_ values that rely on only the linear portion of the sigmoidal dose-response (Figure S2B) and the biphasic portion of the curve to calculate AUCs.

We detected significant differences in PPQ susceptibilities strongly impacted by *pfcrt* genotype for all three traits (Figure 2), with *p* values equal to 2.2e-13 for PSA, 1.0e-08 for AUC and 1.0e-06 for LP-IC_50_ (Figure 2A). Additionally, QTL mapping of the PSA phenotype revealed a highly significant 144 kb region on chr 7 (LOD score of 10.3) that centered on *pfcrt* (Figure 3A). This strong peak is responsible for 62% of the observed inherited phenotypic variance in the PSA phenotype across the progeny set, whereas inheritance of this allele is responsible for 48% of the variance in the distribution of the AUC phenotype. QTL mapping of AUC and LP-IC_50_ identify single QTLs on this chr 7 locus (LOD scores of 6.9 and 5.2 respectively) (Figure 3C,D).

**Figure 2:**
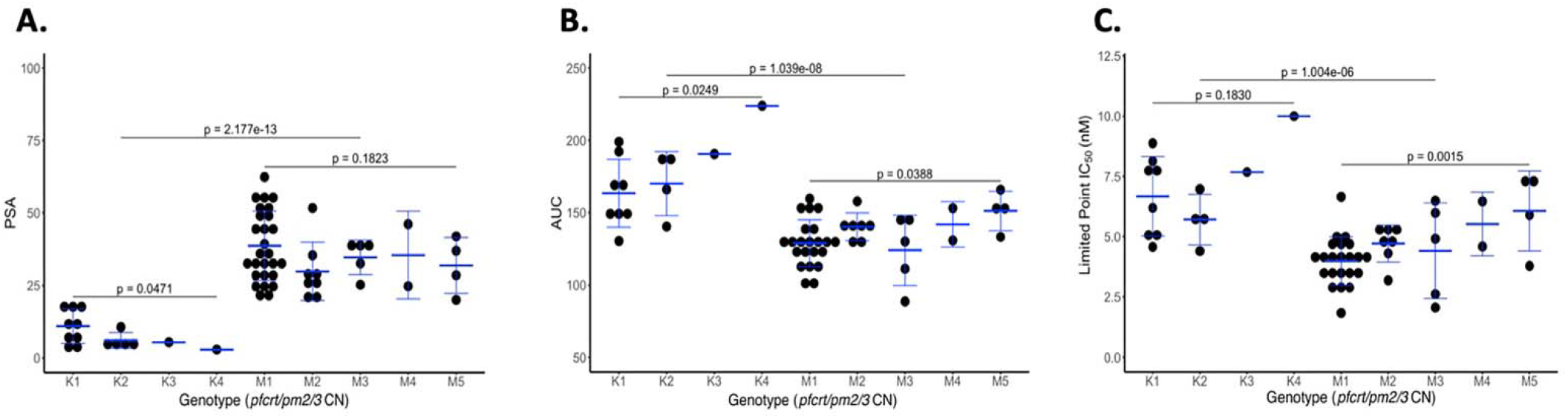
PfCRT mutations determine the majority of the variation in PPQ response. Progeny of the KH004 × Mal31 cross were grouped based on inheritance of parental allele at pfcrt (KH004 or Mal31) and pm2/3 copy number and were phenotyped ing 3 different measurements of resistance: PSA (A), AUC (B), and LP-IC_50_ (C). We find that with all 3 phenotypes, inheritance the KH004 pfcrt allele leads to significantly decreased susceptibility to PPQ. Additionally, we observed that increased copies pm2/3 led to additional decreases in susceptibility, however the impact on CNV with respect to pfcrt varied by phenotype.

**Figure 3:**
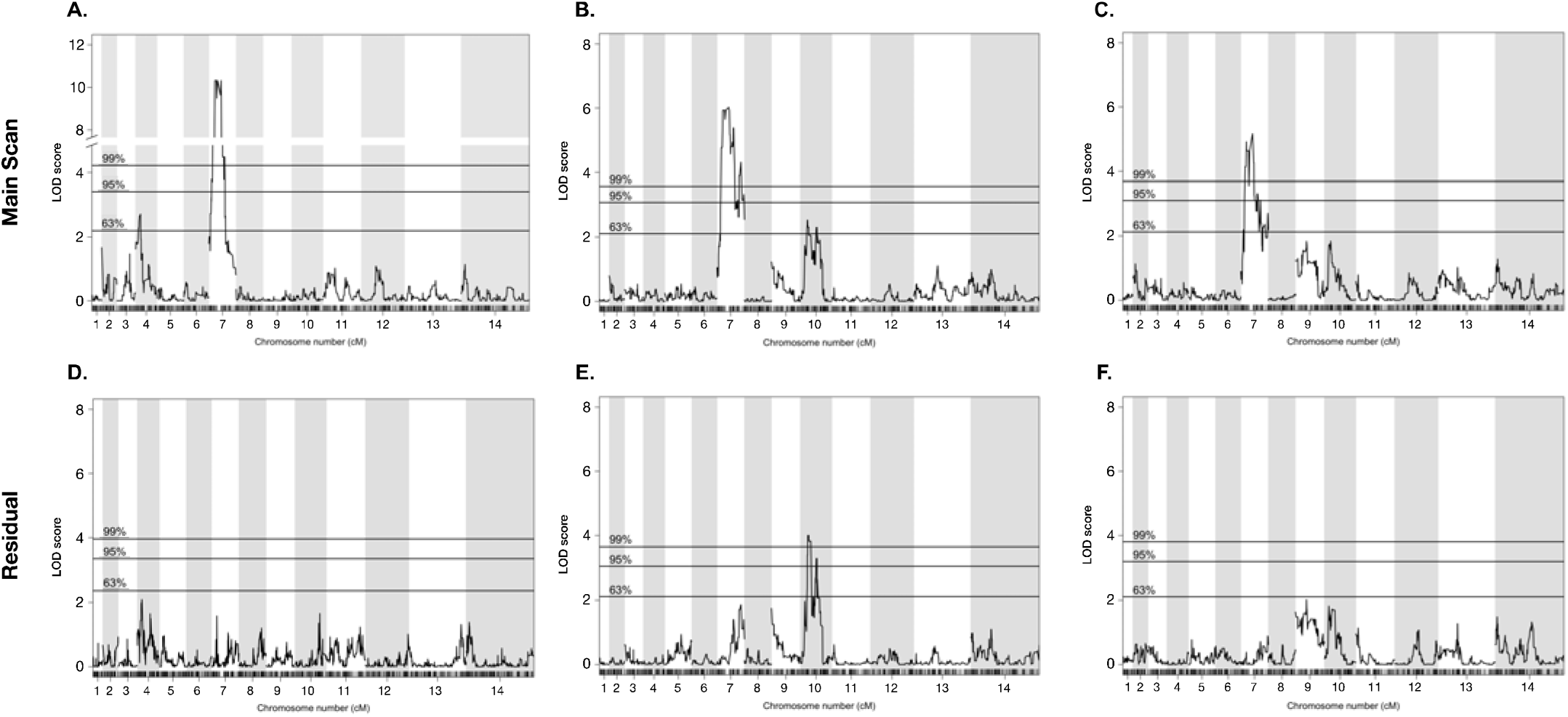
QTL mapping identifies major and secondary loci associated with PPQ-R. We performed QTL mapping based on asite response to PPQ using our 3 metrics of resistance: PSA, AUC, and LP-IC_50_. For each of the 3 phenotypes, a region of 7 centered around pfcrt was most strongly associated with the variation associated with each phenotype. This further ports that pfcrt is the major factor associated with decreased susceptibility. To eliminate the potentially masking effect of the 7 QTL, residual scans of variation were performed. These scans identified a secondary QTL on chr 10 which was associated AUC. Pairing QTL mapping with resistance phenotyping further supports the strong influence on pfcrt as well as identifies ondary contributors to PPQ-R.

### pm2/3 CNV contributes epistatically to increased PPQ resistance

On top of the major segregation of PPQ responsiveness based on *pfcrt* inheritance, we further detected that the trait distributions within *pfcrt* genotype sub-clusters associate with *pm2/3* CNV (Figure 2). However, unlike the progeny distribution shaped by the *pfcrt* allele across all three phenotypes, the effect of *pm2/3* amplification varied depending on the *pfcrt* allele and by phenotype measurement. For example, for PSA, we see an increase in the degree of PPQ resistance associated with increased *pm2/3* CN only in parasites carrying the KH004 *pfcrt* allele (Figure 2A).

To examine the role of *pm2/3* CN independent of other loci, we utilized two sets of parasites: isogenic (IBD) progeny and selfed parental parasites, are identical genotypes except for differences in CN. These parasites are PPQ-S because they inherited the Mal31 *pfcrt* allele. Notably, we did not recover isogenic progeny with the KH004 *pfcrt* allele but displaying different CN. For both PSA and AUC, amplification from single copy *pm2/3* to 4 copies in a PPQ-R (KH004 *pfcrt*) background significantly impacts both phenotypes, however, variation of the number of multiple copies did not significantly impact either phenotype in individuals expressing the Mal31 *pfcrt* allele (Figure 4).

**Figure 4:**
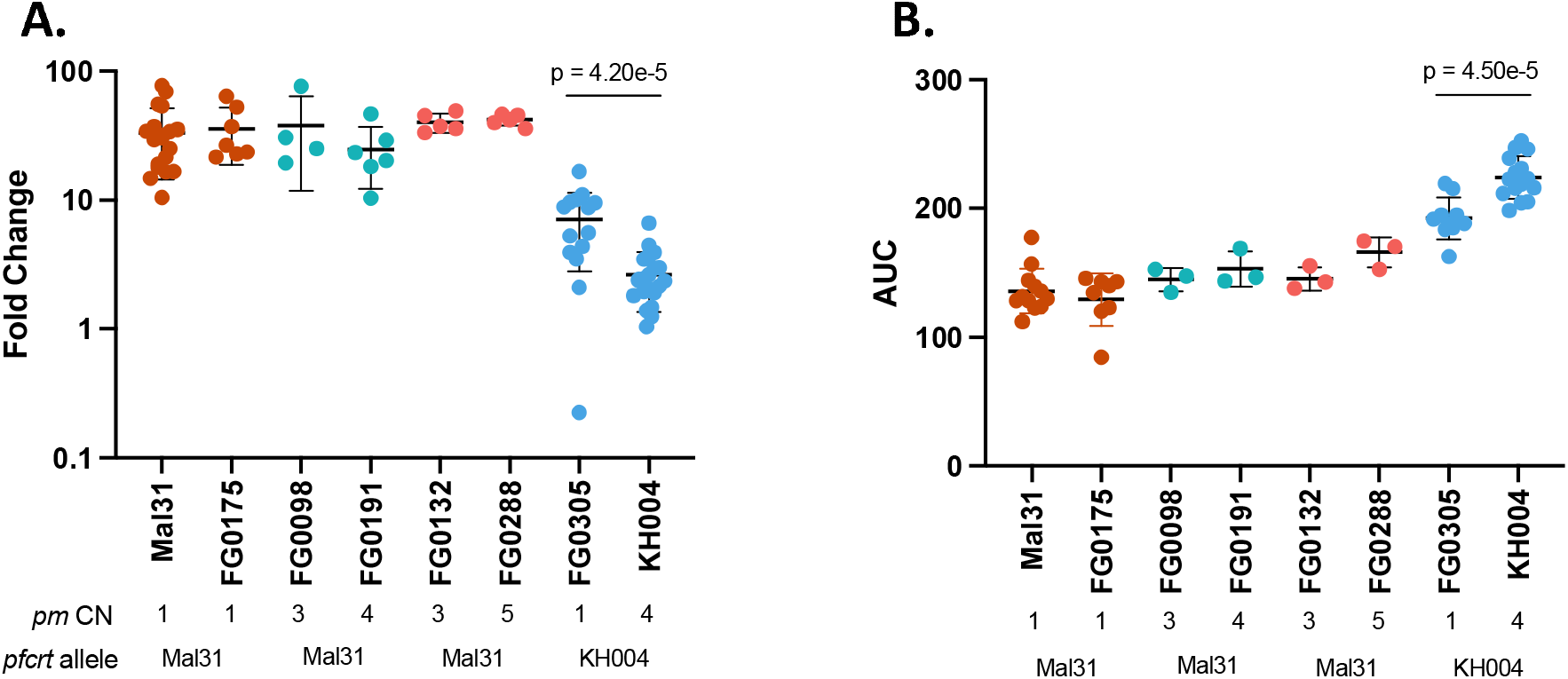
E**f**fect **of plasmepsin II/III copy number on PPQ-R is dependent on genetic background.** Four groups of isogenic parasites which differ only in pm CN were assayed using A) PSA and B) AUC to identify the isolated role of pm CNV independent of genetic background. Of which, FG0175 is a selfed Mal31 progeny; FG0098 and FG0191 were IBD; FG0132 and FG0288 were IBD; FG0305 is a selfed KH004 progeny. We found that CNV in a Mal31-pfcrt background does not significantly impact either phenotype.

### Identification of additional loci contributing to PPQ response using QTL mapping

Due to the large and potentially masking effect of *pfcrt*, secondary scans of residual variation were examined for all traits after statistically removing the effect from that chr 7 locus. While no secondary effects were identified for either the PSA or LP-IC_50_ phenotypes, residual scans of the AUC phenotype identify a significant (LOD = 4.0) QTL on chr 10, which contains 61 genes (95% CI) (Table S5). Additionally, we used a two-dimensional, two-QTL genome scan to identify and characterize significant interactions between loci that contribute to the various resistance phenotypes (Table 1). Using a two-locus model, we identified a significant additive interaction between loci on chr 7 and chr 10 (LOD additive: 10.7; 5% threshold) for the AUC phenotype. Conversely, no QTL interactions were identified using the PSA phenotype.

**Table 1:** Additive interactions influencing PPQ-R phenotypes. Two-dimensional genome scan identifies interacting loci on chromosomes 7 and 10 contribute to the AUC phenotype . We have identified that these two loci interact additively in their contributions to AUC. We identified no interacting loci associated with PSA phenotype. Significant LOD scores are bolded and have surpassed the 5% threshold.

| Phenotype | Pairs of Loci | Position locus 1 | Position locus 2 | LOD interaction (epistasis) | LOD additive |
| --- | --- | --- | --- | --- | --- |
| AUC | Chr 7: Chr 10 | 357253 | 485200 | 1.39 | **10.7** |

**Table 2:** Prevalence of G367C mutation in SEA. Based on available data from MalariaGEN Pf7, this novel polymorphism has en observed throughout SEA, although at low levels, since 2012. Apart from two samples, this mutation is found in PPQ-R mples and is also accompanied by increased copies of pm2/3. ^a^ Two samples from Vietnam carried a single pm2/3 copy and was deemed sensitive independent of pfcrt polymorphisms. ^*^ PPQ-R-associated PfCRT substitutions include: T93S, H97Y, F145I, I218F, M343L, G353V. The C350R substitution was omitted as it arose independently out of a 7G8 background and has not been identified on top of a Dd2 background.

| Country | % of multicopy *pm2/3* samples | # of G367C samples | # of documented PPQ-R mutations* |
| --- | --- | --- | --- |
| Cambodia (n = 1267) | 42.1% | 12 | 327 |
| Vietnam (n= 1404) | 39.0% | 14<sup>a</sup> | 676 |
| Laos (n= 991) | 16.4% | 3 | 169 |
| Global population (n = 16203) | 7.90% | 29 | 1204 |

### PfCRT G367C likely confers PPQ resistance

Comprehensive analysis of KH004 × Mal31progeny points to *pfcrt* as the major determinant of PPQ resistance, along with modulatory effects from other loci. Inheritance of the *pfcrt* allele from KH004 was the major factor shared by PPQ-R parasites (Figures 2 and 4). KH004 carries a Dd2-like *pfcrt*, differing from Dd2 by only a G367C substitution. While the Dd2 *pfcrt* haplotype confers resistance to CQ and other antimalarials, it is not associated with PPQ resistance (25, 27–29). Thus, we suspect the G367C substitution is the major determinant of PPQ-R in KH004. We further explored PPQ susceptibility among the parental strain, KH004, by studying a selfed KH004 parasite (FG0305) that carries only one copy of *pm2/3*), Dd*2,* and NHP4026 (recently isolated from parent at Thailand-Myanmar border (45) that carries a Dd2 *pfcrt* allele and one copy of *pm* 2/3). PSA shows clearly that both parasites (parental KH004 and selfed KH004) with G367C are significantly PPQ-R, while those without G367C (*Dd*2 and NHP4026) have PPQ-S PSA values similar to the African parasites NF54 and Mal31, which have a 3D7-like *pfcrt* (Figure S4).

Reported drug resistance-associated mutations in PfCRT generally reside in one of the 10 transmembrane (TM) domains and, specifically, in regions that make up the central cavity of the transporter (33, 46). The G367C substitution is distinct from experimentally validated PPQ-R-associated mutations in that it resides in the digestive vacuole (DV)-exposed portion of TM9 and is not part of the core structure of the transporter’s central cavity (27, 28, 33, 47, 48) (Figure S3).

This novel G367C mutation was noted by Ross et al (25) to be present in SEA at low prevalence in a small geographic focus. Because the KEL1/PLA1 lineage has continued to spread throughout SEA coinciding with the continuing emergence of PPQ-R, we analyzed the prevalence over time of this novel mutation using the MalariaGEN Pf7 dataset. Of 16,203 global *pfcrt* haplotypes, 29 samples carried G367C (Table 3), first identified in 2012 and restricted to Cambodia but has more recently emerged in neighboring countries Vietnam and Laos (Table S6). Furthermore, 93% of these 29 clinical samples also carry multiple copies of *pm2/3*. We also note the timeline of the emergence of these novel PPQ-R conferring *pfcrt* mutations: Kelch13- mediated ART-R mutations emerged first while the rise of novel PPQ-R-associated *pfcrt* mutations quickly followed the increase of samples expressing multiple copies of *pm2/3* (Figure S5).

### Impacts of pfcrt and pm2/3 inheritance on parasite blood-stage fitness

Our data across multiple resistance phenotypes support that *pfcrt* is the primary driver of PPQ-R, with a boost to the degree of resistance resulting from increased copies of *pm2/3.* It has also been well-documented that mutations in major resistance-associated genes can be associated with a loss in parasite fitness (30, 49, 50). Deleterious fitness effects of the KH004 *pfcrt* allele are clearly evident from the reduced representation of this allele in the progeny recovered from the genetic cross (Figure S1). For these novel resistance-associated genotypes to increase in global prevalence in the absence of selection from widespread drug treatment, parasites must preserve fitness either through low-cost resistance mutations or the acquisition of secondary compensatory mutations that preserve or improve fitness. We conducted competitive growth assays between progeny of the KH004 × Mal31 cross and both parental parasites to assess blood-stage fitness differentials (Figure S6). Co-cultures were maintained for a maximum of 40 d or until one parasite reached fixation (>90% of culture), with samples being collected every 2 d. Progeny vs. parent competitions revealed that most progeny had a relative fitness situated between two parents, while four progeny parasites were outside of the parental fitness range (two more fit than KH004 and two less fit than Mal31), with no significant clustering in fitness based on inherited *pfcrt* and *pm2/3* genotypes (Figure S6).

Based on the similar fitness phenotypes generated from competitions against the parental parasites, a select subset of 18 progeny and both parents were set up in overlapping competitions against common competitors, resulting in 212 competitive outcomes (win, lose, or draw). The outcomes of these pairwise competitions were used to generate an Elo Rating for each parasite using the *EloRating* package in R (51). Elo ratings provide a weighted ranking of parasite fitness based on competitive growth outcome as well as the fitness ranking of the competitor, with a higher Elo Rating representing a greater competitive fitness level. Elo Ratings for these 20 parasites (parents plus progeny) provide a much finer resolution of fitness differences between parasites of seemingly similar fitness. Unlike the competitions against both parents, Elo Ratings provide a more continuous phenotype, but still do not cluster based on *pfcrt* and *pm2/3* genotype (Figure 5A). However, QTL analysis using Elo phenotypes reveals a highly significant (1% threshold) on chromosome 9 (LOD score of 4.5), which encompasses a 35 kb region containing 15 genes (Figure 5B). Parasites with the greatest competitive fitness were associated with inheritance of this 35 kb region from the KH004 parent. Sequence comparison between KH004 and Mal31 for these 15 genes identified 4 non-synonymous coding SNPs in two genes: PF3D7_0934100 and PF3D7_0934700 (Table S7).

**Figure 5:**
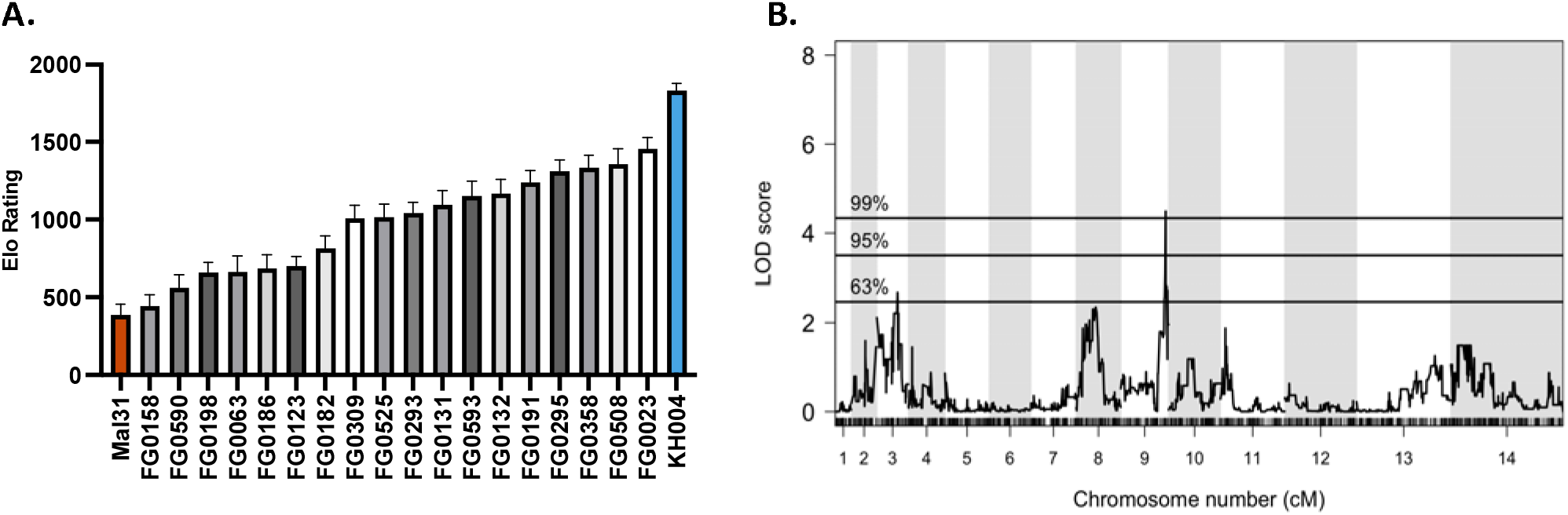
Q**T**L **mapping of fitness-associated loci using Elo rating phenotype. A)** Mean Elo rating of 18 progeny + 2 parents of the KH004 × Mal31 genetic cross based on 50 permutations of the order of competitions. Elo rating was calculated using the R package EloRating and was based on the each parasite’s competitive growth assay outcome. Phenotype distribution is ordered from lowest Elo (least fit) on the left to highest Elo (most fit) on the right. **B)** QTL scan based on Elo phenotype reveals a highly significant peak (1% threshold) on chromosome 9. This peak encompasses a 35,264 bp region which contains 15 genes. Genes identified from this QTL scan are proposed to directly contribute to parasite fitness.

## Discussion

In this study, we used a genetic cross between a multidrug-resistant Cambodian parasite and a drug-sensitive isolate from Malawi to characterize the genetic contributors to PPQ-R. Previous work on this subject has relied on naturally resistant parasites isolated from patience or *in vitro* experimental manipulations to test specific mutations’ ability to confer PPQ-R (13, 16, 20, 25). Use of a genetic cross allows us to track the inheritance and impact on phenotypes of alleles previously associated with resistance, such as SNPs in *pfcrt* or amplification of *pm2/3*, while also searching for additional trait-impacting loci across the genome . This classical genetic approach demonstrates that the inheritance of the KH004 PfCRT (Dd2 + G367C) is the main driver of PPQ-R, with the amplification of *pm2/3* serving an additive role amongst resistant parasites.

The KH004 parent of this cross expresses a novel G367C substitution in PfCRT, previously noted in SEA but not associated with PPQ-R. Through comparisons between KH004, Dd2, and other SEA parasites, we observed a > 30-fold change in PSA between KH004 and parasites expressing a standard Dd2-like PfCRT, indicating a direct connection between this substitution and PPQ-R. While the G367C substitution has not previously been connected to PPQ-R, this residue is within the binding site of the structurally similar chloroquine derivative perfluorophenylazido biotinylated chloroquine (AzBCQ) (52). This finding of an association of this residue with PPQ-R contributes to a broader understanding of PfCRT structure that influences a wide range of drug susceptibilities, i.e. the range of *pfcrt* haplotypes shaped by evolution to drug pressure. The major *pfcrt* haplotypes associated with PPQ-R have all evolved on a 7G8 or Dd2 background, and transfection of these mutations into different *pfcrt* backgrounds generates slight decreases in PPQ-R levels compared to those in a Dd2 background (28, 33).

Previously studied PPQ-R-associated PfCRT substitutions including T93S, H97Y, C101F, F145I, M343L, and G353V were suspected due to their increasing prevalence in sites using the DHA+PPQ (25, 33). The novel G367C substitution expressed by KH004 arose in western Cambodia in 2012 and was found at a low abundance in the population. Since its initial emergence, the Dd2+G367C haplotype has continued to persist at a consistently low abundance but has spread to neighboring countries, Laos, and Vietnam, with the latter accounting for the majority of identifications in recent years. As most of SEA has transitioned from DHA+PPQ to alternative combination therapies as frontline treatments, Vietnam continues to use PPQ, which might explain the recent increase in abundance of this novel haplotype within the country (1, 53, 54). On the other hand, the introduction of this haplotype into Laos can potentially be accounted for by a recent introduction of PPQ-R parasites into the country from Cambodia (55).

Due to unusual PPQ-parasite interactions that are not effectively captured by standard dose-response (IC_50_) curves, we utilize three unique phenotypes to illuminate different aspects of PPQ-R: AUC, limited point IC_50_, and PSA. By using these different assays, we deconstruct the resistance phenotype into three related but distinct biological readouts, survival under high single dose PPQ (PSA), low dose growth inhibition (LP-IC_50_), and shape of the dose-response curve under high PPQ concentrations (AUC), to refine our search for the genetic factors controlling these phenotypes (13, 18, 21, 22). QTL Mapping for each trait identified mutant *pfcrt* as the main underlying contributor to resistance. Additionally, statistically removing the effect of the *pfcrt* locus allowed for the identification of secondary contributors to the corresponding phenotype. While the PSA and LP-IC_50_ residual scans did not identify any secondary loci, an additional locus on chr 10 had a highly significant association with the AUC phenotype in the absence of *pfcrt* (Figure 2). One of the genes with the strongest association within this region is *autophagy-related gene 18* (*atg18*). KH004 contains a single non-synonymous SNP in *atg18*, leading to a Thr38Ile substitution, whereas Mal31 carries the WT 3D7-like variant of this gene, highlighting a candidate mutation for further investigation. This protein localizes to the parasite DV and has been associated with decreased susceptibility to several antimalarials such as DHA, artemether (AM), and PPQ, while also being connected to increased survival under nutrient deprivation (56–58). Due to its previous association with PPQ-R and the association only with the AUC phenotype, *atg18* may contribute to the non-traditional dose-response curve encountered when studying *in vitro* PPQ-R. Due to the fact that inheritance of the KH004 *pm2/3* allele does not directly correspond to a set number of copies, the physical map used for QTL analysis only associates allelic inheritance, not copy number, with each phenotype. Given that *pm2/3* CN is what impacts PPQ susceptibility, we do not observe any QTL encompassing this locus on either or main or residual scans.

Previous studies hypothesized that instead of directly modulating PPQ-R, expressing multiple copies of *pm2/3* amplification may play a compensatory role in parasite fitness (22). This hypothesis, paired with extensive previous findings on the fitness costs of *pfcrt* resistance mutations, led us to assess the competitive fitness levels of our progeny with various combinations of *pfcrt* and *pm2/3* (29, 47, 49). By comparing progeny fitness to KH004 and Mal31 parents, there found no significant clustering of *pfcrt/pm2/3* genotype combinations that suggest a simple genetic determination of parent-derived fitness. Consequently, we leveraged all-on-all competitions to generate Our novel implementation of Elo rankings can efficiently compare head-to-head fitness levels across a large progeny set to generate a refined phenotype for QTL mapping. Using this approach, we identified a highly significant association between parasite fitness and a narrow 35 kb region on chr 9. This region is includes genes involved in development, transcriptional regulation, and metabolism including two prioritized candidate genes with coding variations between KH004 and Mal31. Neither gene has been linked previously to parasite fitness.

Deeper knowledge of the genetic determinants of PPQ resistance can point to novel control and prevention strategies. Our classical genetic approach informed our understanding of the major drivers of resistance, including a novel *pfcrt* mutation, as well as identified secondary genetic factors that contribute to parasite resistance and fitness. Incorporation of a range of PPQ-R phenotype measures into our analysis revealed unique genetic regions and their interactions associated with changes in each phenotype. By combining this novel genetic cross with rigorous *in vitro* resistance and fitness phenotyping, we have identified the genetic architecture which underlies decreased susceptibility to PPQ. While the DHA+PPQ ACT continues to be used as a frontline ACT, it is crucial to understand the genetic underpinnings of resistance.

---

**Figure S1:**
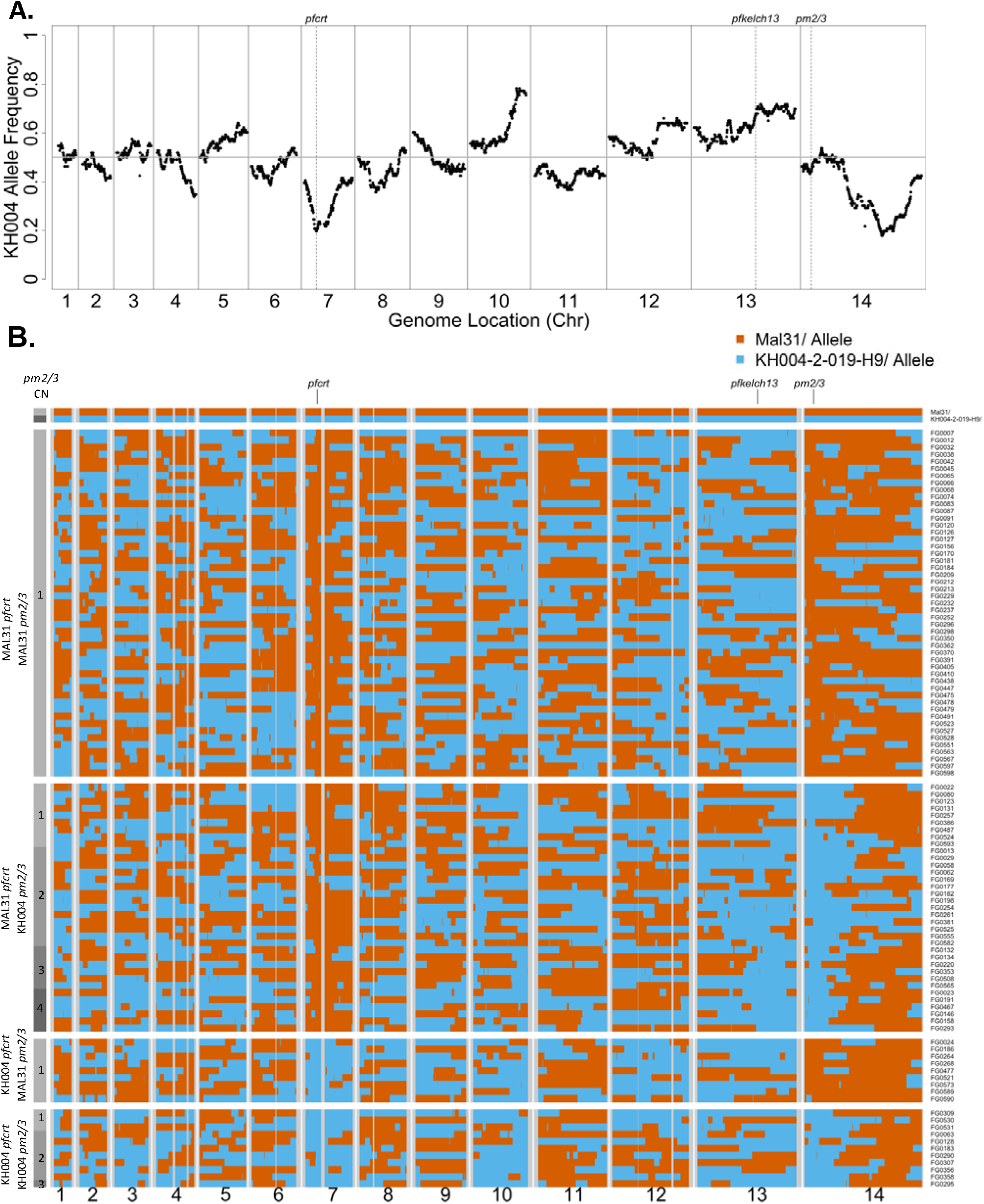
Inheritance breakdown of KH004xMal31 genetic cross. A) From this cross, we see two strong skews in inheritance on chromosomes 7 and 14, both favoring the Mal31 allele. This selective inheritance has previously been observed in other genetic crosses between Southeast Asian and African parasites. The chromosome 7 peak is centered around pfcrt and therefore potentially represents a fitness cost associated with the inheritance of the KH004 allele at these positions. B) Aside from the selective inheritance on chromosomes 7 and 14, we observe no significant co-inheritance patterns between pfcrt, pm2/3, or kelch13. Inheritance of the KH004 allele is denoted in blue and the Mal31 allele in orange.

**Figure S2:**
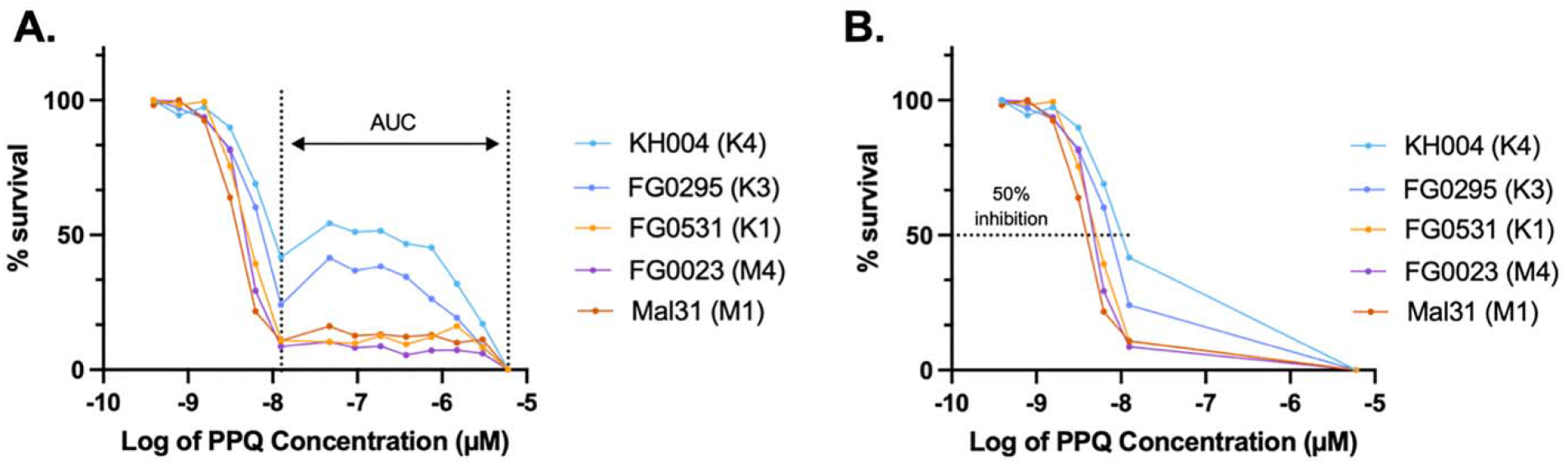
Shape of dose-response curve changes based on pfcrt and pm2/3 genotype. Traditional IC_50_ measurements have been a challenge with PPQ due to the biphasic dose-response curve observed in PPQ-R parasites. A) We have found inheritance of the KH004 pfcrt allele and multiple copies of pm2/3 are required for producing a biphasic curve. Progeny inheriting the Mal31 pfcrt allele produce do not produce a biphasic curve, regardless of CNV. Additionally, all parasites expressing a single copy of pm2/3 produce a sigmoidal curve. To account for these various dose-response curve shapes, AUC is used as a measure of resistance, with a greater AUC indicating decreased PPQ susceptibility. B) By using a limited number of points in our dose response curve (LP-IC_50_), we can eliminate the secondary peak in the curve and measure an IC50 independent of original curve shape.

**Figure S3:**
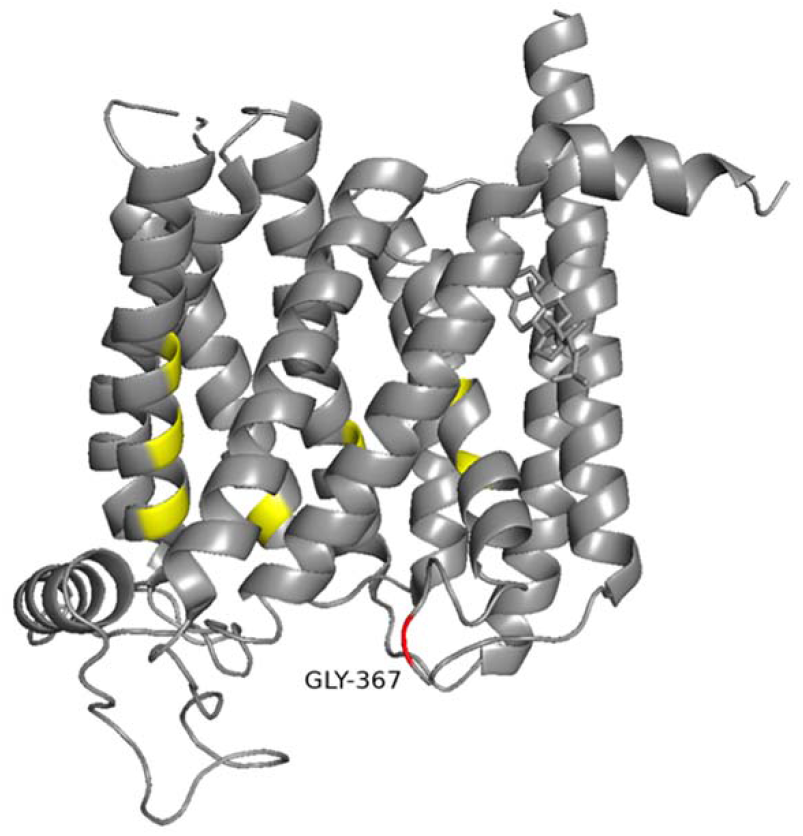
Structural position of G367C Substitution: A) Alphafold model of Dd2 PfCRT with previously documented in vitro PPQ-R associated mutations identified (yellow). Additional highlighted AA substitutions and the novel G367C substitution carried by KH004 (red).

**Figure S4:**
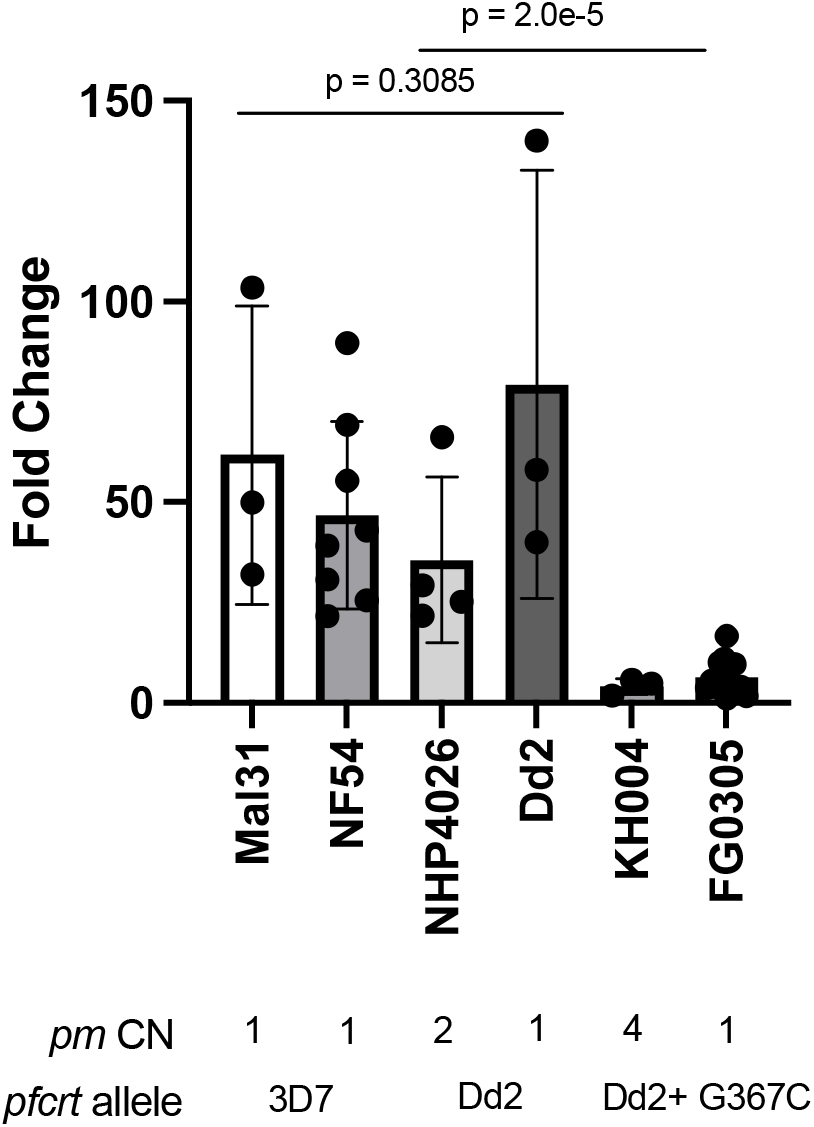
PfCRT G367C directly confers PPQ-resistance. PSA between KH004, FG0305 (selfed KH004 with single copy pm) and two other southeast Asian parasites (Dd2 and NH4026) which carry a Dd2-like PfCRT allele, and two African parasites carrying a WT PfCRT identifies that a Dd2 PfCRT background alone is not sufficient for PPQ-R. This further supports the conclusion that PfCRT G367C directly confers PPQ resistance.

**Figure S5:**
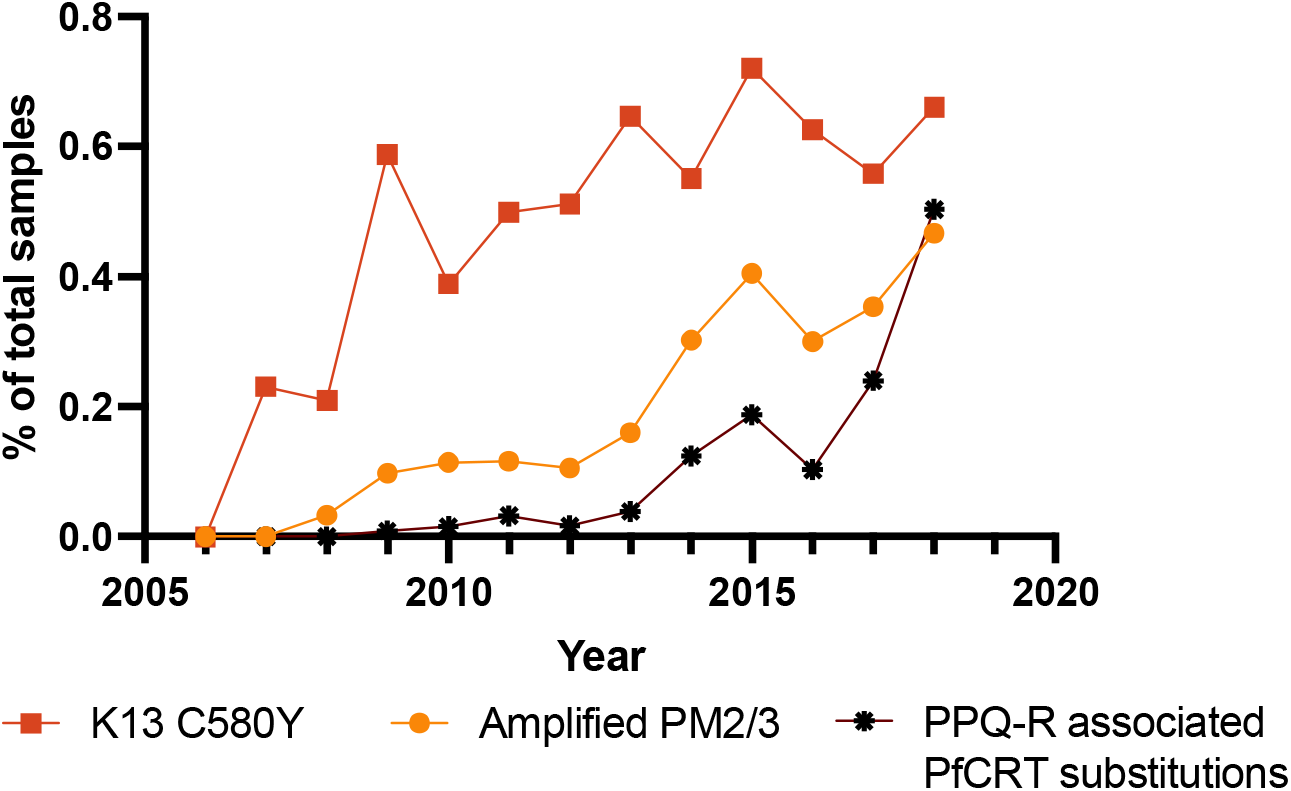
Rise of pm2/3 amplification and novel pfcrt mutations associated with PPQ-R in the GMS. Using available data from MalariaGEN Pf7 release (n = 3,359), samples from Cambodia, Vietnam, and Laos were analyzed for the association between pm2/3 amplification and novel PPQ-R conferring pfcrt mutations. The rise in novel pfcrt PPQ-R mutations immediately following amplification of pm2/3 further confirms the important roles of both pm2/3 and pfcrt in the evolution of PPQ-R.

**Figure S6:**
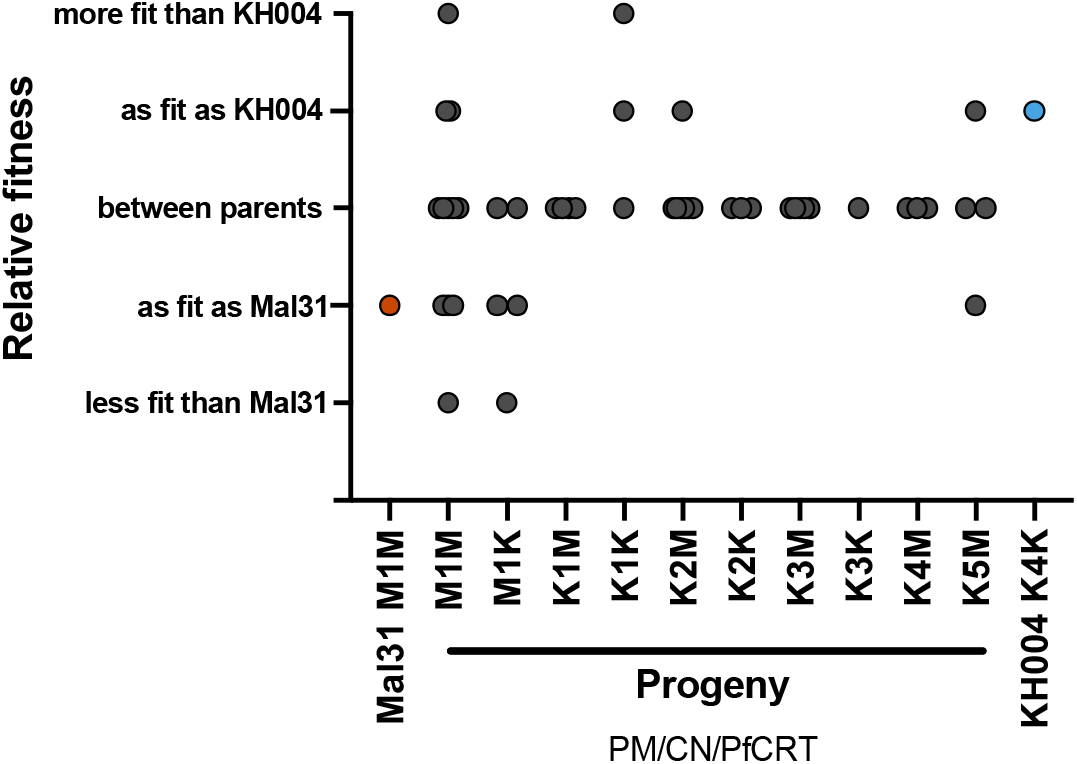
Competitive growth assessment between progeny and parents. Head-to-head competitive growth assays were set up between progeny and parents and were ranked based on competitive outcome. The fitness level of most progeny fell in between the two parents and no apparent clustering was observed between relative fitness and pfcrt or pm2/3 genotype.

## Methods

### Ethics approval and consent to participate

The study was performed in strict accordance with the recommendations in the Guide for the Care and Use of Laboratory Animals of the National Institutes of Health (NIH), USA. To this end, the Seattle Children’s Research Institute (SCRI) has an Assurance from the Public Health Service (PHS) through the Office of Laboratory Animal Welfare (OLAW) for work approved by its Institutional Animal Care and Use Committee (IACUC). All of the work carried out in this study was specifically reviewed and approved by the SCRI IACUC.

### Preparation of the genetic crosses

We generated the crosses using FRG NOD huHep mice with human chimeric livers and *A. stephensi* mosquitoes as described by Vaughan *et al.* (*41*). Two individual recombinant pools were generated for each cross, by using different cages of infected mosquitoes. To start each cross, gametocytes from both parental parasite strains were diluted to 0.5% gametocytemia in a human serum erythrocyte mix, to generate infectious blood meals (IBMs). IBMs from each parent were mixed at equal proportions and fed to three cages of mosquitos (150 per cage).

We examined the mosquito infection rate and oocyst number per infected mosquito 7-10 days post-feeding. Fifteen mosquitoes were randomly picked from each cage and dissected under microscopy. Sporozoites were isolated from infected mosquito salivary glands and 2-4 million sporozoites from each cage of mosquitoes were injected into three FRG huHep mice (one cage per mouse), intravenously. To allow the liver stage-to-blood stage transition, mice are infused with human erythrocytes six and seven days after sporozoite injection. Four hours after the second infusion, the mice are euthanized and exsanguinated to isolate the circulating ring stage *P. falciparum*-infected human erythrocytes. The parasites from each mouse constitute the initial recombinant pools of recombinant progeny for genetic mapping experiments. We maintained the initial pools in AlbuMAX supplemented RPMI media; we genome sequenced aliquots from each pool to check allele frequencies from both parents.

### Library preparation and sequencing

We used Qiagen DNA mini kit to extract and purify the genomic DNA, and Quant-iT™ PicoGreen® Assay (Invitrogen) to quantify the amount of DNA. For samples with less than 50ng DNA obtained, whole genome amplification (WGA) was performed before NGS library preparation. WGA reactions were performed following Nair et al (Nair et al., 2014). Each 25 μl reaction contained at least 5ng of *Plasmodium* DNA, 1× BSA (New England Biolabs), 1 mM dNTPs (New England Biolabs), 3.5 μM of Phi29 Random Hexamer Primer, 1× Phi29 reaction buffer (New England Biolabs), and 15 units of Phi29 polymerase (New England Biolabs). We used a PCR machine (SimpliAmp, Applied Biosystems) programmed to run a “stepdown” protocol: 35 °C for 10 min, 34 °C for 10 min, 33 °C for 10 min, 32 °C for 10 min, 31 °C for 10 min, 30 °C for 6 h then heating at 65 °C for 10 min to inactivate the enzymes prior to cooling to 4 °C. Samples were cleaned with AMPure XP Beads (Beckman Coulter) at a 1:1 ratio. We constructed next generation sequencing (NGS) libraries using 50-100 ng DNA or WGA product following the KAPA HyperPlus Kit protocol with 3-cycle of PCR. All libraries were sequenced at 150bp pair-end using Illumina Novaseq S4 or Hiseq X sequencers. We sequenced all bulk samples to a minimum coverage of 100×.

### Mapping and genotyping

We individually mapped whole-genome sequencing reads for each library against the *P. falciparum* 3D7 reference genome (PlasmoDB, release32) using the alignment algorithm BWA mem (http://bio-bwa.sourceforge.net/) under the default parameters. The resulting alignments were then converted to SAM format, sorted to BAM format, and deduplicated using picard tools v2.0.1 (http://broadinstitute.github.io/picard/). We used Genome Analysis Toolkit GATK v3.7 (https://software.broadinstitute.org/gatk/) to recalibrate the base quality score based on a set of verified known variants (42).

After alignment, we excluded the highly variable genome regions (subtelomeric repeats, hypervariable regions and centromeres) and only performed genotype calling in the 21 Mb core genome (defined in (42)). We called variants for each sample using HaplotypeCaller, and calls from every 100 samples were merged using CombineGVCFs with default parameters. Variants were further called at all sample-level using GenotypeGVCFs, with parameters: -- max_alternate_alleles 6 --variant_index_type LINEAR --variant_index_parameter 128000 -- sample_ploidy 2 -nt 20. We further filtered the variants calls by calculating the recalibrated variant quality scores (VQSR) of genotypes from parental parasites. Loci with VQSR less than 1 or not distinguishable between two parents were removed from further analysis. The variants in VCF format were annotated for predicted functional effect on genes and proteins using snpEff v4.3 (https://pcingola.github.io/SnpEff/) with 3D7 (PlasmoDB, release32) as the reference.

### Cloned progeny analysis

To identify unsuccessfully cloned progeny, we measured the multiplicity of progeny samples with F_WS_ (59), which is configured in moimix (https://github.com/bahlolab/moimix). Samples with F_WS_ < 0.9 were assumed to be non-clonal and were removed from further analysis. Allele frequencies across the genome were also plotted and manual inspected to detect further possible mix infections.

We calculated Identity-by-descent (IBD) between clonal progeny and parents with hmmIBD (60) under default parameters. The proportions of shared IBD were used to determine relatedness among parental and progeny parasites and to identify genome regions inherited from each parent: 1) progeny with >90% shared IBD with either of the parents were assumed to result from selfing; 2) progeny with <90% shared IBD with both parents were defined as recombinants; 3) recombinant progeny with >90% IBD against each other were defined as non-unique clones (61).

### Nanopore sequencing of parental parasites

We used an optimized a Phenol:Chloroform DNA isolation protocol that generates 500 ng to 3 µg large molecular weight genomic DNA from 40 mL of in vitro blood cultures. In brief, (i) short, fragmented DNA (<60kb) is removed using Small Read Eliminator Kit (Nanopore); (ii) after DNA clean-up, we prepared sequencing libraries with Ligation Sequencing Kit (Nanopore), which adds barcode to each sample. We used the Nanopore MinION Mk1C sequencer to generate long reads. We obtained 3.5 Gb (150× genome coverage) and 2.5 Gb (105×) of long-read sequencing data for Mal31 and KH004. Fifty percent of the data obtained comprise reads with length >62Kb (N50 = 62kb) with the longest read of 485 kb.

We mapped the nanopore long reads to *plasmepsin* gene sequences using *blastn* (https://blast.ncbi.nlm.nih.gov/Blast.cgi). Reads that cover all three *plasmepsin* genes and >10kb flanking regions on both sides were then extracted and compared to 3D7 reference genome to identify the repeat unit at *plasmepsin* genes.

### Plasmepsin II/III copy number identification

We analyzed mapped reads on the 67 kb extended loci of the *plasmepsin* genes (from 260,000 to 327,000 at chromosome 14), which including 28 kb upstream *plasmepsin* I and 28 kb downstream *plasmepsin* III. The number of reads mapped onto each position (coverage) were determined from the deduplicated BAM file using *bedtools* (https://bedtools.readthedocs.io/en/latest/). We visualized reads mapping to the 67 kb region using the Integrative Genomics Viewer (IGV) (Fig S1A). The repeat unit of at *plasmepsin* genes was confirmed by both IGV plot and nanopore long-read sequences. We calculated the *Plasmepsin* copy number using mean[coverage at repeat unit]/mean[coverage at extension regions].

### Linkage Analysis

Linkage between the *pfcrt*, *pm2/3* and *pfkelch13* loci was assessed by calculating D and D′ and utilizing Fisher’s exact test to determine significance. Association between *pfcrt* or *pfkelch13* alleles and *pm2/3* CN were assessed by binning multicopy *pm2/3* CN and conducting Fisher’s exact test.

### Parasites used in this study

This study utilizes parents and progeny of the KH004 × Mal31 genetic cross generated by utilizing human liver-chimeric. From this cross, one hundred and four unique recombinant progeny were recovered through limited dilution cloning at 0.3 cells per well. Individual wells containing parasites were identified through qPCR and analysis of progeny relatedness and allele inheritance were based on the methods outlined in Button-Simons et al. (45). Methods for sequencing of progeny, construction of the physical map, and generation of visual recombination map are all based on Button-Simons et al. (45). Analysis of allelic co-inheritance utilized the entire progeny set of one hundred and four parasites and phenotyping assays utilized fifty unique parasites, forty-eight progeny and the two parental parasites.

### Parasite culture

Cryopreserved *P. falciparum* stocks of KH004 × Mal31 progeny and KH004-020-019-H9 and Mal31-9040-C11 parents were thawed and cultured in complete media (CM) consisting of 0.5% Albumax II (Gibco, Life Technologies) supplemented RPMI 1640 with L-glutamine (Gibco, Life Technologies) with additional 50 mg/L hypoxanthine (Calbiochem, Sigma-Aldrich), 25 mM HEPES (Corning, VWR), 10ug/ml gentamycin (Gibco, Life Technologies), and 0.225% sodium bicarbonate (Corning, VWR). Parasite cultures were maintained at 5% hematocrit in O+ red blood cells (RBC) (Interstate Blood Bank, Memphis, TN) in separate flasks and maintained consistent temperature (37°C) and atmosphere (5% CO_2_/5% O_2_/90% N_2_). Parasitemia of cultures was kept below 2% and media changes were performed every 48 hours, corresponding with one intraerythrocytic development cycle.

### PfCRT Structure Analysis

Three-dimensional homology model of PfCRT was predicted using AlphaFold (62, 63) and accessed through the PDB database (6UKJ) (33). Additionally, the F′(ab) fragment used in 7G8 PfCRT cryo-EM structure elucidation (33) was removed from the structure. Protein mutagenesis and visualization of resistance-associated mutations were conducted using PyMol software (v2.5.4; Schrödinger, LLC).

### Drug susceptibility assays

Prior to susceptibility assay setup, parasites were synchronized using a single layer of 70% Percoll (Sigma Aldrich) in 1X RPMI with 13.3% sorbitol in phosphate buffered saline. 400µL of packed erythrocytes infected by a majority of schizont stage parasites were resuspended in 2mL of incomplete media, layered over the Percoll layer, and centrifuged (1561 x g for 10 in, no brake). The top layer was removed to isolate late-stage schizonts, washed twice with incomplete media, resuspended at 5% hematocrit in complete media, and placed on a shaker for 4 hours at 37°C. Following the 4 hour incubation, parasitemia and stage were determined through flow cytometry by staining with SYBR Green I and SYTO 61 and analyzed on a Guava easyCyte HT (Luminex) after 50,000 events were counted. If cultures were >70% ring-stage, they were diluted to 0.15% parasitemia at 2% hematocrit and set up in a 96-well plate at a volume of 150µL per well. Within each assay plate, two technical replicates of each parasite were exposed to 10 concentrations of a 2-fold dilution series of PPQ as well as both untreated parasite and uninfected RBC controls. Parasites were exposed to PPQ for 72 hours. Parasite density was determined using SYBR Green and dose-response curves were generated by plotting percent survival against the log of drug concentration. Analysis of IC_50_ and AUC was performed using Prism 9.0 (GraphPad). A nonlinear log inhibitor vs response 4 parameter dose-response curve was generated based on percent survival. To generate the limited point curve, data points which made up the biphasic portion of the plot were excluded to allow for a sigmoidal shaped curve, which was then used for IC_50_ calculations.

### PSA

PSA was set up as originally described (18) with synchronized early ring stage parasites being expose to a single 200nM dose of PPQ for 48 hours. This protocol was modified to resemble the extended recovery RSA (eRRSA) phenotype (64), in which samples are analyzed at 120 hours post-exposure instead of 72 hours. This extended timepoint has been shown to provide superior differentiation between resistant and sensitive parasites and allows for finer resolution between PPQ-S parasites. Parasite prevalence was measured through qPCR and survival was calculated as fold change between untreated and treated samples for three technical replicates (3 replicates within the plate as well as 3 independent qPCR reads) and at least 3 biological replicates per sample. Additionally, to ensure consistency between our resistance phenotypes, parasites from the same synchronization were used for both PSA and IC_50_/AUC susceptibility assays.

### Competitive growth assays

To determine relative fitness levels between parasites, we used competitive growth assays between progeny of the KH004 × Mal31 cross and both parental parasites as a proxy for *in vitro* fitness. Parasitemia and stage of parasites was quantified using flow cytometry 15 hours after synchronization with a 70% Percoll gradient. Synchronized parasites were each adjusted to 0.5% parasitemia at 5% hematocrit and set up in a 1:1 ratio (1% total parasitemia) in 96-well plates as previously described in (37, 39). Parasitemia was assessed (Giemsa-stained slides and microscopy) every two days, in accordance with one intraerythrocytic life cycle, adjusted to 1% parasitemia and supplemented with fresh blood and media. Samples from each well were collected at regular intervals upon sample dilution and stored at -80°C for genotyping. These competitive growth assays were maintained for a maximum of 40 days, or until one parasite reached fixation (>95% of the co-culture). Parasites were deemed to have equivalent fitness levels if by 40 days neither parasite reached fixation.

To analyze relative parasite densities in each competition, six microsatellite markers (MS) were used for genotyping (Table S8) to differentiate parasites in co-culture. PCR amplification was performed using the Phusion Blood Direct PCR Kit (ThermoFisher, cat #F547L) for 30 cycles as previously described in (37). Amplified microsatellites were analyzed using the Applied Biosystems 3730xI DNA Analyzer (ThermoFisher). Raw fragment analysis data was uploaded to the Thermo Fisher Connect platform and analyzed using the microsatellite analysis tool. The proportion of the two competing parasites in co-culture was determined by taking the ratio of fluorescent peak height associated with the corresponding PCR product size.

Due to the large number of competitive growth assays required to directly compete all parasites against each other, all parasites were competed against both parents. Additionally, a genetically diverse subset of parasites were competed against each other. Based on the outcomes of the progeny v progeny competitions, an Elo rating system, which has previously been used for animal dominance hierarchies, was utilized to assign a quantitative score to each parasite based on their win/loss record while also taking into account the record of the parasite competed against (65). The package “EloRating” (51) in R Studio (2023.03.0+386). was used for assigning scores. In order to reduce the effect of competition order on ranking, the competitive outcomes file was randomized 1000 times to generate 1000 Elo scores which were then averaged. The distribution of Elo ratings figure was created using Prism 9.0 (GraphPad).

### QTL Analysis

Statistical analysis of the QTL data for this cross was performed using the computational methods previously described (66). Blood stages of *P. falciparum* are haploid, therefore, only two genetic classes are present for each locus. QTL mapping was performed using the “R/qtl” package (67) in R Studio. One thousand permutations of the trait values determined the 37%, 5%, and 1% genome-wide significance thresholds, and the strength of each linkage was expressed as a logarithm of the odds (LOD) score (68). The main QTL and corresponding mean trait values from these scans were used to obtain estimates of residual empirical thresholds to identify secondary loci that contribute to the phenotype of interest (68). After identifying QTLs associated with each phenotype, QTL interaction was tested using the scantwo function of the “R/qtl” package and was based on 1,000 permutations.

### Global haplotype analysis

Analysis of global *pfcrt* haplotypes was performed using publicly available sequences through the Pf7 dataset (MalariaGEN) (12). PPQ resistance status for this dataset was inferred based on *plasmepsin II/III* amplification, therefore, all samples labeled as PPQ-R are assumed to express more than one copy of these genes.

### Data Availability

All data needed to evaluate the conclusions in the paper are present in the paper and/or the Supplementary Materials. All raw sequencing data have been submitted to the NABI Sequence Read Archive (SRA, https://www.ncbi.nlm.nih.gov/sra) under the project number of PRJNA524855. Additional data related to this paper may be requested from the authors.

### Code Availability

The code used in analysis and data analysed are available at GitHub through the following links: https://github.com/kbuttons/MalariaGeneticCrosses-MAL31xKH004

## Acknowledgments

This work was supported by National Institutes of Health (NIH) program project grant P01 AI127338 (to MF) and by NIH grant R37 AI048071 (to TJCA). Nanopore sequencing was supported by Texas Biomedical Research Institute Forum Grant 20-04866 (to XL). Work at Texas Biomedical Research Institute was conducted in facilities constructed with support from Research Facilities Improvement Program grant C06 RR013556 from the National Center for Research Resources. The parental line, Mal31, used in the *Mal31×KH004* cross was sampled from a Malawian patient in 2016 as part of a cross-sectional study funded by the Wellcome Trust of Great Britain (Grant no. 099992/Z/12/Z to SCN). We thank the patients who provided parasites used in this work.

## Additional Information

Additional figures and tables are available for this paper.

